# Prediction of Post Traumatic Epilepsy using MRI-based Imaging Markers

**DOI:** 10.1101/2024.01.12.575454

**Authors:** Haleh Akrami, Wenhui Cui, Paul E. Kim, Christianne N. Heck, Andrei Irimia, Karim Jebri, Dileep Nair, Richard M. Leahy, Anand A. Joshi

## Abstract

Post-traumatic Epilepsy (PTE) is a debilitating neurological disorder that develops after traumatic brain injury (TBI). Despite the high prevalence of PTE, current methods for predicting its occurrence remain limited. In this study, we aimed to identify imaging-based markers for the prediction of PTE using machine learning. Specifically, we examined three imaging features: lesion volumes and resting-state fMRI-based measures of functional connectivity and amplitude of low-frequency fluctuation (ALFF). We employed three machine learning methods, namely, kernel support vector machine (KSVM), random forest, and a neural network, to develop predictive models. Our results showed that the KSVM classifier, with all three feature types as input, achieved the best prediction accuracy of 0.78 AUC (Area Under the Receiver Operating Characteristic (ROC) curve) using nested cross-validation. Furthermore, we performed voxel-wise and lobe-wise group difference analyses to investigate the specific brain regions and features that the model found to be most helpful in distinguishing PTE from non-PTE populations. Our statistical analysis uncovered significant differences in bilateral temporal lobes and cerebellum between PTE and non-PTE groups. Overall, our findings demonstrate the complementary prognostic value of MR-based markers in PTE prediction and provide new insights into the underlying structural and functional alterations associated with PTE.

## 1. Introduction

Traumatic Brain Injury (TBI) survivors often carry a tremendous burden of disability as a result of their injuries [1]. Such injuries can have wide-ranging physical and psychological effects with some signs or symptoms that appear immediately after the traumatic event, while others appear days or weeks later. TBI is one of the major causes of epilepsy [2] yet the link between TBI and epilepsy is not fully understood. Post-traumatic epilepsy (PTE) refers to recurrent and unprovoked post-traumatic seizures occurring after 1 week [3]. Patients with PTE perform worse across several clinical and performance metrics such as independence and cognitive scores and have a significantly reduced quality of life [4]. They are prone to higher rates of mental illness such as depression and addiction [5]. Significant risk factors for the development of seizures > 1 week after TBI include seizures within the first week, acute intracerebral hematoma (especially subdural hematoma), brain contusion, increased injury severity, and age > 65 years at the time of injury [6]. The incidence of PTE ranges from 4% to 53%, with the risk approaching 50% in cases with direct injury to the brain parenchyma. PTE risk also varies with the time after injury, age range under study, as well as the spectrum of severity of the inciting injuries [6, 7, 8]. Epidemiological studies have found that PTE accounts for 10%–20% of symptomatic epilepsies and 5% of all epilepsies [9, 10]. It is estimated that in the USA and the European Union (EU), with a total population of about 800 million, at least 0.5 million surviving individuals live with PTE [3]. Data on the economic burden of PTE are unavailable, but some idea is provided by the lifetime cost of TBI on average, which in the USA is around $200,000 per case scaled to 2004 prices [11, 12]. Thus, in addition to the personal burden, the economic burden caused by PTE is also substantial. Therefore, prediction and if possible, prevention of PTE remains an important challenge.

Biomarkers for post-traumatic epilepsy can vary from imaging and electrophysiologic measurements to changes in gene expression and metabolites in blood or tissues. MRI imaging offers a non-invasive, radiation-free powerful modality for marker development. While a wealth of biomarkers exist when the epilepsy condition is already established, these markers can only reveal mechanisms that exist after the epileptogenic process, which can allow partial or full pharmacoresistance to establish before treatment starts [13, 8, 14]. Prognostic markers and approaches to identify the risk of post-traumatic epilepsy would eliminate the need to wait for spontaneous epileptic seizures to occur before starting treatment. The ability to identify high-risk subjects can enable the mitigation of risks to subjects whose seizures could result in serious injury or death.

The pathogenesis of spontaneous recurrent seizures are certainly multifactorial. Once established, seizure threshold, which is a measure of the balance between excitatory (glutaminergic) and inhibitory (GABA-ergic) forces in the brain [15, 16], is thought to vary over time depending on several factors such as periodicities in seizure occurrence [17]. Current anti-seizure medications raise the seizure threshold and thus reduce the propensity for seizures to occur. An individualized prognostic marker for the development of PTE could be used in clinical trials to study compounds which may have true anti-epileptic potential. Current medications used for epilepsy only treat the symptoms not the underlying pathophysiology which leads to epilepsy. The value of individualized prognostic markers is five-fold [18]: (i) Prediction of the development of an epilepsy condition: prognostic markers that eliminate the need to wait for spontaneous epileptic seizures to occur would reduce the time and cost required for TBI patients to start participation in clinical trials, and also the risks to subjects whose seizures could result in serious injury or death, (ii) Identification of the presence and severity of tissue capable of generating spontaneous seizures: an imaging-based marker can identify anomalous brain regions, which could help in surgical as well as noninvasive treatment planning even before the condition is established. (iii) Measuring progression after the condition is established: MR imaging-based markers can help in quantifying the progression of epilepsy and understanding pharmacoresistance. Identification of localized biomarkers of epileptogenic brain abnormalities would allow longitudinal tracking of seizure threshold at later time points and would presumably reveal the time points when the epileptogenic process reaches a critical point so that clinical seizures would likely occur. (iv) Creating animal models of PTE: The identified markers can be used to create animal models, and prediction algorithms can be used for more cost-effective screening of animal models for treatment with potential anti-epileptogenic and antiseizure drugs and devices. (v) Cost reduction in clinical trials by screening patients: PTE risk prediction can be used for recruitment into clinical trials for potential anti-epileptogenic interventions by enriching the trial population with identified patients that are at higher risk for developing epilepsy.

Prediction and if possible, prevention of the development of PTE is a major unmet challenge. Animal studies in adult male Sprague-Dawley rats have shown the potential of using MRI-based image analysis for finding biomarkers for PTE [19, 20]. These studies point towards the involvement of the perilesional cortex, hippocampus, and temporal lobe in PTE [10]. Despite important progress, brain imaging is still underexploited in the context of PTE biomarker research. Numerous human neuroimaging studies have provided important insights into TBI [21, 22, 23] and epilepsy [24, 25, 26], but imaging-based PTE prediction work is scarce.

Group analyses of TBI patients compared to controls revealed volume reductions in brain tissue across multiple cortical areas including the corpus callosum, corona radiata; anterior, mid, and posterior cingulate gyrus, precuneus, and parahippocampal gyrus [22, 27, 23, 28, 29], as well as subcortical regions, including the hippocampus, amygdala, putamen, globus pallidus, caudate, and midbrain [21, 22, 27, 28, 30, 31].

Clinical and research studies in epilepsy often include both anatomical (MRI, CT) and functional (PET, EEG, MEG, ECoG, depth electrodes, fMRI) mapping. While epileptogenic zones can be found in almost any location in the brain, the temporal lobe and the hippocampus are the most common sites causing focal epileptic seizures [26]. Multimodal MRI and PET imaging has been used to predict the laterality of temporal lobe epilepsy [32, 26]. Extensive changes in brain networks due to epilepsy were reported using PET, fMRI, and diffusion imaging [24, 20, 32, 33, 26]. Research using resting-state fMRI (rs-fMRI) over the last two decades has uncovered important properties of brain dynamics and network organization by revealing the existence of patterns of spontaneous neural activity that occur in the absence of a specific task or stimulus. These patterns, known as resting-state networks, have been found to be consistent across individuals and are thought to reflect the underlying functional organization of the brain [34]. The presence of lesions in TBI patients is expected to alter these resting-state brain dynamics either locally, through changes in the lesioned area, or at the level of networks affected by the lesion [35]. In addition, previous work has shown that cases of focal epilepsy are often associated with changes in network activity extending beyond the seizure onset zone [36, 37]. Interestingly, Zhou et al. [38] trained an SVM classifier on brain imaging data for the diagnosis of mesial temporal lobe epilepsy, and found that combining fMRI and structural MRI features provided better classification than either modality alone. Taken together, these studies motivate the exploration of local and network-level brain abnormalities as potential predictive markers of PTE in patients who have suffered from a TBI.

A few recent studies have employed machine learning to identify functional brain changes that may serve as PTE biomarkers [39, 33, 40]. Common features used in such brain-based prediction approaches are the pair-wise correlation patterns observed between rs-fMRI signals. However, because of the high dimensionality of such connectivity metrics, using them as features in a machine learning framework can quickly lead to model overfitting. A standard approach to dealing with this issue is to use a dimensionality reduction method such as principal components analysis (PCA) to reduce the dimension of the feature space to a subset of principal components that represents nevertheless most of the information in the data. Moreover, regularized models [41, 42] with a ridge or lasso penalty can also be used to select a subset of features and prevent overfitting by penalizing the weights. In the specific case of exploiting brain connectivity features in a data set with limited number of subjects, the presence of groups of highly correlated features leads to an ill-conditioned feature space. As a result, methods that use simple penalties that discard most of the correlated features can become unstable [42].

The goal of the present study was to probe the utility of multiple structural and functional features extracted from MR imaging in characterizing PTE, as well as predicting its occurrence. We explored both classical group-level statistics and cross-validated machine learning methods. Our results extend previous reports by uncovering key MR-based differences between TBI patients who develop PTE and those who do not. In addition, we were able to leverage machine learning analyses to assess the relative contribution of different types of structural and functional features to the out-of-sample prediction of PTE.

## 2. Materials and Methods

### 2.1. Data

We extracted functional and structural features from two datasets:

1. The Maryland TBI MagNeTs dataset [43]: Of the 113 individual data sets, 72 (36 PTE and 36 non-PTE groups) were used for group-level difference comparisons as well as for supervised training and testing of a machine learning algorithm (i.e. constructing a model from training samples to predict the presence of PTE in previously unseen data). The remaining 41 non-PTE subjects (total 113) were used to train an artificial neural network for automatic lesion delineation, using a method recently developed by our group [44].
2. The TRACK-TBI Pilot dataset [45]: 97 subjects from this data set, in addition to the 41 non-PTE subjects from MagNeTS, were used to train the artificial neural network for automated delineation of lesions as described briefly below in Section 2.2.2. A more detailed description can be found in [44].

#### 2.1.1. Maryland MagNeTs data

This is our main dataset for group comparisons as well as for PTE prediction. The dataset was collected as a part of a prospective study that includes longitudinal imaging and behavioral data from TBI patients with Glasgow Coma Scores (GCS) in the range of 3-15 (mild to severe TBI). Injury mechanisms included falls, bicycle or sports accidents, motor vehicle collisions, and assaults. The individual or group-wise GCS, injury mechanisms, and clinical information is not shared. The imaging data are available from FITBIR (https://fitbir.nih.gov), with FLAIR, *T*_1_, *T*_2_, fMRI, diffusion, and other modalities available for download. In this study, we used imaging data acquired within 10 days after injury, and seizure information was recorded using follow-up appointment questionnaires. Exclusion criteria included a history of white matter disease or neurodegenerative disorders including multiple sclerosis, Huntington’s disease, Alzheimer’s disease, Pick’s disease, and a history of stroke or brain tumors.

The imaging was performed on a 3T Siemens TIM Trio scanner (Siemens Medical Solutions, Erlangen, Germany) using a 12-channel receiver-only head coil. For statistical analysis, we used 36 fMRI subjects with PTE (25M/11F) from this dataset and 36 randomly selected fMRI subjects without PTE (28M/8F)[43, 46]. The age range for the epilepsy group was 19-65 years (yrs) and 18-70 yrs for the non-epilepsy group. Our analysis of population differences was performed using *T*_1_-weighted, *T*_2_-weighted, and FLAIR MRI as well as resting fMRI [43]. The remaining 41 subjects with TBI but without PTE were used for training the automatic lesion detection algorithm. Standard gradient-echo echo-planar resting-state functional MR imaging (repetition time msec/echo time msec, 2000/30; flip angle, 75°; field of view, 220 × 220 mm; matrix, 128 × 128; 153 volumes) was performed in the axial plane, parallel to a line through the anterior and posterior commissures (section thickness, 5 mm; section gap, 1 mm) and positioned to cover the entire cerebrum (spatial resolution, 1.72 × 1.72 × 6.00 mm) with an acquisition time of 5 minutes 6 seconds. The individuals were instructed to close their eyes for better relaxation but to stay awake during the imaging protocol.

#### 2.1.2. TRACK-TBI Pilot dataset

This is a multi-site study with data across the injury spectrum, along with CT/MRI imaging, blood biospecimens, and detailed clinical outcomes [45]. Here we use 3T MRI data in addition to information collected according to the 26 core Common Data Elements (CDEs) standard developed by the TrackTBI Neuroimaging Working Group. The 3T MRI protocols (implemented on systems from General Electric, Phillips, and Siemens) complemented those used in the Alzheimer’s Disease Neuroimaging Initiative (ADNI) with *T*_*R*_/*T*_*E*_ = 2200/2.96 ms, an effective *T*_*I*_ of 880 ms, an echo spacing time of 7.1 ms, a bandwidth 240*Hz*/*pixel*, and a total scan time of 4 minutes and 56 seconds. The data is available for download from https://fitbir.nih.gov

To train the unsupervised deep learning model, a variational autoencoder for lesion delineation as described in Section 2.2.2, we used 2D slices of brain MRIs from a combined group of 41 TBI subjects (33M/8F, age range 18-82 yrs) from the Maryland TBI MagNeTs study [43] and 97 TBI subjects (70M/27F, age range 11-73 yrs) from the TRACK-TBI Pilot study [45]. These TBI data were taken from patients without PTE, and are strictly distinct from the set of 72 subjects which we subsequently used for statistical testing and PTE prediction.

### 2.2. Methods

#### 2.2.1. Preprocessing

Pre-processing of the MR datasets was performed using the BrainSuite software (https://brainsuite.org). The three modalities (*T*_1_, *T*_2_, FLAIR) were coregistered with each other by registering *T*_2_ and FLAIR to *T*_1_, and the result was co-registered to the MNI atlas (Colin 27 Average Brain) [47] by registering *T*_1_ images to the MNI atlas using a rigid (translation, scaling and rotation) transformation model. As a result, all three image modalities were registered to a common MNI space at 1 mm^3^ resolution. Skull and other non-brain tissue were removed using BrainSuite [48], and brain extraction was performed by stripping away the skull, scalp, and any non-brain tissue from the image. This was followed by tissue classification and generation of the inner and pial cortex surfaces. Subsequently, for training and validation of the lesion detection model, all images were reshaped into 128 × 128 pixel images and histogram-equalized to a lesion-free subject.

The extracted cortical surface representations and brain image volumes for each subject were jointly registered to the BCI-DNI Brain Atlas (http://brainsuite.org/svreg_atlas_description/) [49] using BrainSuite’s Surface-Volume Registration (SVReg18a) module [50, 51]. The BCI-DNI brain atlas is from a single subject with parcellation defined by anatomical landmarks. SVReg uses anatomical information from both the surface and volume of the brain for accurate automated co-registration, which allows consistent surface and volume mapping to a labeled atlas. This co-registration establishes a one-to-one correspondence between individual subjects’ *T*_1_ MRIs and the BCI-DNI brain atlas. The deformation map between the subject and the atlas encodes the deformation field that transforms images between the subject and the atlas.

We used the BrainSuite fMRI Pipeline (BFP) to process the rs-fMRI subject data and generated grayordinate representations of the preprocessed rs-fMRI signals [52]. BFP is a software workflow that processes fMRI and T1-weighted MR data using a combination of software that includes BrainSuite, AFNI, FSL, and MATLAB scripts to produce processed fMRI data represented in a common grayordinate system that contains both cortical surface vertices and subcortical volume voxels. Starting from raw T1 and fMRI images, BFP produces processed fMRI data co-registered with BrainSuite’s BCI-DNI atlas and includes both volumetric and grayordinate representations of the data.

#### 2.2.2. MRI Based Measures: Lesion Detection

To extract lesions from the anatomical MRI data we used an unsupervised framework which we recently developed to automatically detect lesions in MR data. The method, which has been validated on other data sets, is based on a variational auto-encoder (VAE), a class of auto encoders where the latent representation can be used as a generative model [53, 54, 55]. By training the VAE using nominally healthy (lesion-free) imaging data, the network learns to encode normal brain images. As a result, applying such a model to an image that contains a lesion will yield a VAE-decoded image that does not contain anomalies: the lesions can then be identified from the differences between original and VAE-decoded images. One complication here is that we did not have access to normal imaging data with matching characteristics of the PTE dataset. Instead we trained VAE using the *T*_1_-weighted, *T*_2_-weighted, and FLAIR MRIs in the Maryland TBI MagNeTs dataset [43] leveraging VAEs robustness to outliers [56]. While lesions are present in most of the volumetric TBI images, they are typically confined to a limited region in each brain, so that in any particular anatomical region (at the scale of the major gyri delineated in the BCI-DNI atlas) the fraction of images with lesions is relatively low. In this study the lesions were delineated based on VAE reconstruction error in the FLAIR images. We have previously evaluated performance of this lesion detection algorithm using an independent validation set with delineated lesions [44, 55].

Statistical analyses: Once we determined the lesions using the methods described above, we analyzed the VAE lesion maps using a 1-sided nonparameteric (permutation-based) F-test to determine whether there were any statistically significant differences in the variances of lesion maps across the PTE and non-PTE TBI groups [57]. The null hypothesis is that the variances of lesion maps across subjects in the PTE group is less than or equal to that of the non-PTE group. Our decision to use the F-test was guided by the fact that traumatic brain injuries affect different areas in different subjects across the groups, so that consistently localized differences between the 2 groups were expected to be very unlikely. However, a higher frequency of lesions in a particular regions should result in a higher sample variance in the lesion maps. This rationale was in fact supported by our observation that assessing differences in the group means using a standard t-test did not show any significant effects. In order to apply F-test with permutations, we computed point-wise group variances and computed their ratio to obtain an unpermuted F-statistic. We then permuted the group labels to recompute the F-statistic for *nperm* = 1000 permutations. The *p*-value was computed pointwise by comparing the unpermuted F-statistic to the permuted F-statistics (*nperm* = 1000). Finally, the resulting pointwise map of *p*-values was corrected for multiple comparisons using Benjamini-Hochberg FDR correction [58].

Additionally, we performed a regional analysis by quantifying lesion volume from binarized lesion maps in each ROI using the USCLobes brain atlas [49] (http://brainsuite.org/usclobes-description). The USCLobes atlas segments the brain into larger regions (lobes) than those provided by the default BrainSuite atlases. It has 15 ROIs delineated on the volumetric labels of the atlas: (L/R) Frontal Lobe, (L/R) Parietal Lobe, (L/R) Temporal Lobe, (L/R) Occipital Lobe, (L/R) Insula, (L/R) Cingulate, Brainstem, Cerebellum, and Corpus Callosum.

To identify lesions as binary masks, a one-class SVM [59] was applied to the VAE lesion maps at each voxel and across subjects to identify subjects with abnormally large errors (i.e discrepancies between the original input image and VAE-decoded image) at that voxel [59, 60]. We defined the outliers marked by the one-class SVM as lesions and computed lesion volumes per ROI by counting the number of outlier voxels in each ROI for each subject. In the following we consider this measure to be a proxy for an ROI-wise lesion volume which we use for PTE prediction.

#### 2.2.3. fMRI Based Measures: Connectivity

We compute ROI-wise connectivity using rs-fMRI data and the USCLobes ROIs. The BFP fMRI pipeline produces a standardized fMRI signal in the grayordinate system that is in point-wise correspondence across subjects. Using the USCLobes parcellation with respect to the grayordinates, we computed the ROI-wise signal by averaging over each ROI. A 15x15 matrix was then computed from the Pearson correlations of averaged fMRI signals between each of the 15 ROIs in the USCLobes atlas. We used the elements of the upper-triange of the correlation matrix as a feature vector and applied the Fisher z transform to normalize the feature distribution. This feature vector was subsequently used for classification.

#### 2.2.4. fMRI Based Measures: ALFF

The amplitude of low-frequency fluctuation (ALFF) [61, 62] is an rs-fMRI-based metric that measures the magnitude of spontaneous fluctuations in BOLD-fMRI signal intensity for a given region. We calculated the ALFF metric [62] as the signal power in a frequency band defined by a low- and high-frequency cutoff, which we set to 0.01 Hz and 0.1 Hz, respectively. The ALFF measure was first computed in the native fMRI space and then mapped to the the mid-cortical surface of the USCBrain atlas [49] using the BrainSuite registration method described above.

Statistical analyses: Similar to the statistical assessment of lesion differences, we used a 1-sided F-test (null hypothesis var in PTE <= var in nonPTE populations). We tested for significance using a permutation method (*nperm* = 1000), for voxel-wise group-level comparison of variance across the PTE and non-PTE groups. This resulting *p*-values were corrected for multiple comparisons using the false discovery rate (Benjamini Hochberg procedure) [58]. As a result, we end up with a voxelwise p-value map in the USCBrain atlas space, indicating local differences in slow-frequency fluctuations of BOLD between PTE and non-PTE groups.

#### 2.2.5. PTE prediction Using Machine Learning

In addition to the statistical analyses described above, we examined the same brain features using a supervised machine learning framework[33]. This was motivated by several factors. The machine learning framework allows us to implement an out-of-sample analysis that assess the ability of the features extracted from the data to classify individual subjects. This is particularly important when searching for potential biomarkers. Second, tools such as multi-feature classification and feature importance quantification readily provide useful insights into the significance, complementarity or redundancy across the set of explored features. This is also key when seeking to identify the most efficient prognostic biomarkers. Finally, the diversity of available machine learning algorithms opens novel opportunities to tease apart variable distributions that may be harder to separate using standard statistical tools.

In the machine learning pipeline implemented in the present study, we used the anatomical and functional features described above (i.e. lesion information, connectivity and ALFF) that were extracted from MR imaging data collected during the early (acute) phase prior to onset of PTE. The goal of the data-driven classifier approach is to build a model that learns to distinguish between PTE and non-PTE subjects using labeled training data. We use a leave-one out stratified cross-validation scheme to reduce risks of selection bias and overfitting. To fine-tune model hyperparameters while adhering to a strict separation of training and test data, we used a standard nested cross-validation procedure.

To assess the feasibility of building models that can predict PTE from structural and functional MR data, we implemented a multi-feature binary classification framework using three distinct types of algorithms (details below). We concatenated the extracted features into an input vector, to which we applied PCA to reduce the dimensionality of the feature space [63, 64]. We use the area under the receiver operating characteristic curve (AUC) as the primary performance evaluation metric.

We applied the following three machine learning algorithms:

##### Random Forests [65]

We used a random forest classifier which is an ensemble learning method that works by training multiple decision trees on random subsets of the data and then averaging the predictions of each tree to make a final prediction. This technique reduces overfitting and improves the overall accuracy of the model compared to using a single decision tree. At each iteration, a random subsample of the data is taken, and a new decision tree is fit. This process is repeated multiple times, and the final output is the majority vote of all the decision trees.

##### Support Vector Machines (SVMs) and Kernel-Support Vector Machines (KSVMs)

The basic idea underlying the SVM is to find the hyperplane in a high-dimensional space that maximally separates the different classes [66]. The data points that are closest to the hyperplane are called support vectors and have the greatest impact on the position of the hyperplane. Once the hyperplane is found, new data can be easily classified by determining on which side of the hyperplane they fall. By contrast, a Kernel Support Vector Machine (KSVM), is an extension of the basic SVM algorithm that uses a kernel operator to map the input data into a higher-dimensional space, in which they can be more easily separated. The use of a kernelallows the SVM to handle non-linearly separable data, by finding a higher-dimensional space in which they are linearly separable. One of the most popular kernels for this purpose, and that used here, is the radial basis function (RBF).

##### Multi-Layer Perceptron

We also used a multilayer perceptron (MLP) which is a feedforward artificial neural network where the input is passed through multiple layers of artificial neurons. Each layer applies a non-linear transformation to the input before passingto the next layer [67]. MLPs are trained using back-propagation and stochastic gradient descent. The MLP model we used here consisted of 3 hidden layers (32, 16, and 16 neurons, respectively). While more complex and deeper neural network architectures are available, we chose to use a simple MLP given the limited size of the data at hand (N=72). We expected the RF and SVM algorithms to be more suitable for our classification task, but included the MLP method for the sake of comparison.

## 3. Results

### 3.1. Lesion Analysis

To compare lesion patterns across the PTE and non-PTE patients, we assessed group differences in lesion scores, defined as the difference between the grayscale values in the original anatomical MR images and the VAE-decoded versions thereof (see methods). Statistical assessment using the F-test revealed statistically significant differences (p<0.05, corrected) between the two groups in multiple brain areas (Figure 1). These differences were prominent in the left and right temporal lobes, the right occipital lobe, and the cerebellum, reflecting higher variability of lesion scores in these areas in PTE patients. In addition, our lobe-wise analysis (Table 2) yielded results consistent with voxel-wise analysis, confirming an increased variance in the PTE population relative to non-PTE subjects in the same regions identified in the graordinate-wise analysis.

**Figure 1:**
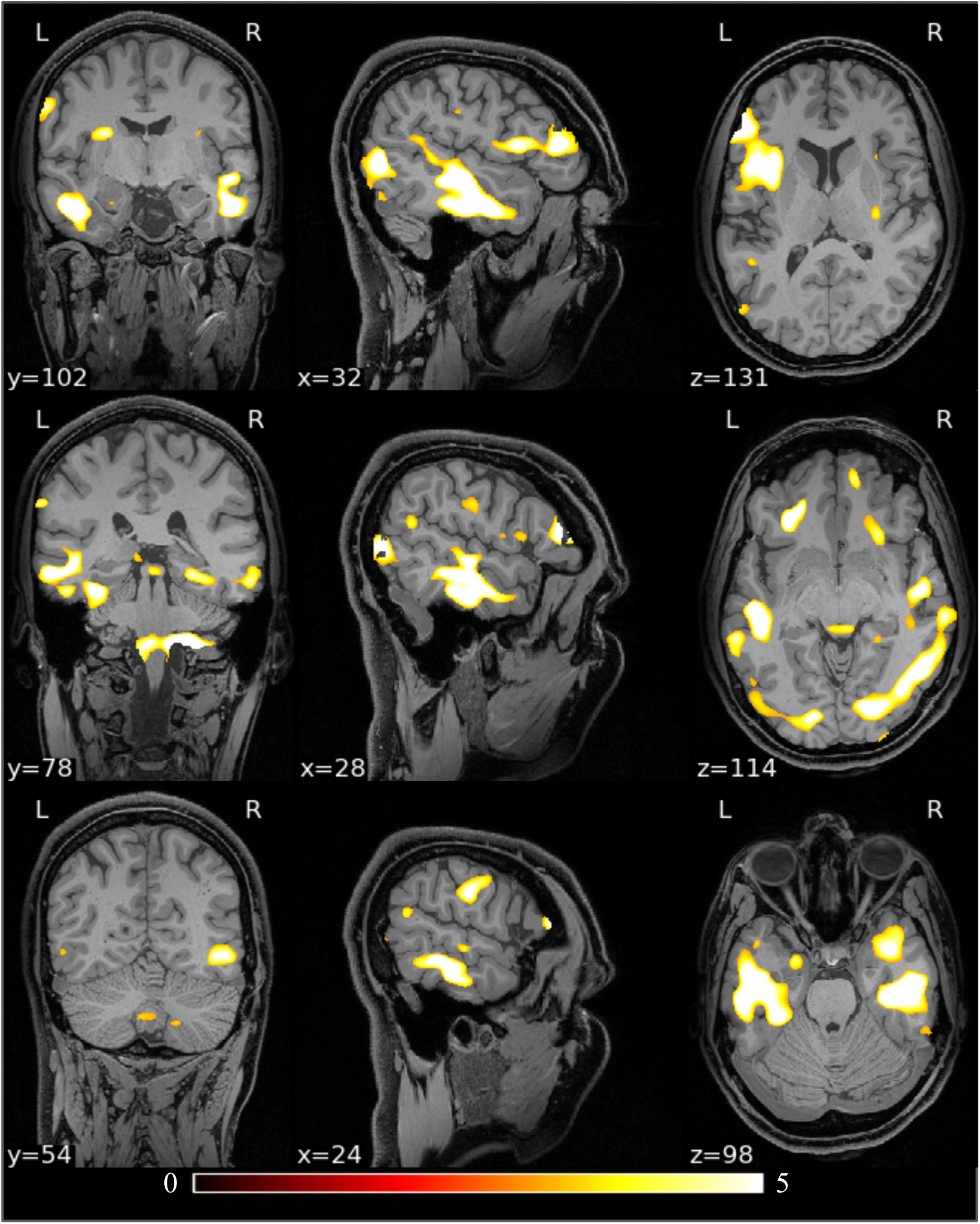
Voxel-based PTE vs. non-PTE group comparison of lesion maps overlaid on the USCBrain atlas. The color code depicts f-values, shown in a region where p-value < 0.05, resulting from the F-test (with permutations). Prominent significant clusters are located in the left temporal lobe, bilateral occipital lobe, cerebellum, and right parietal lobe.

**Figure 2:**
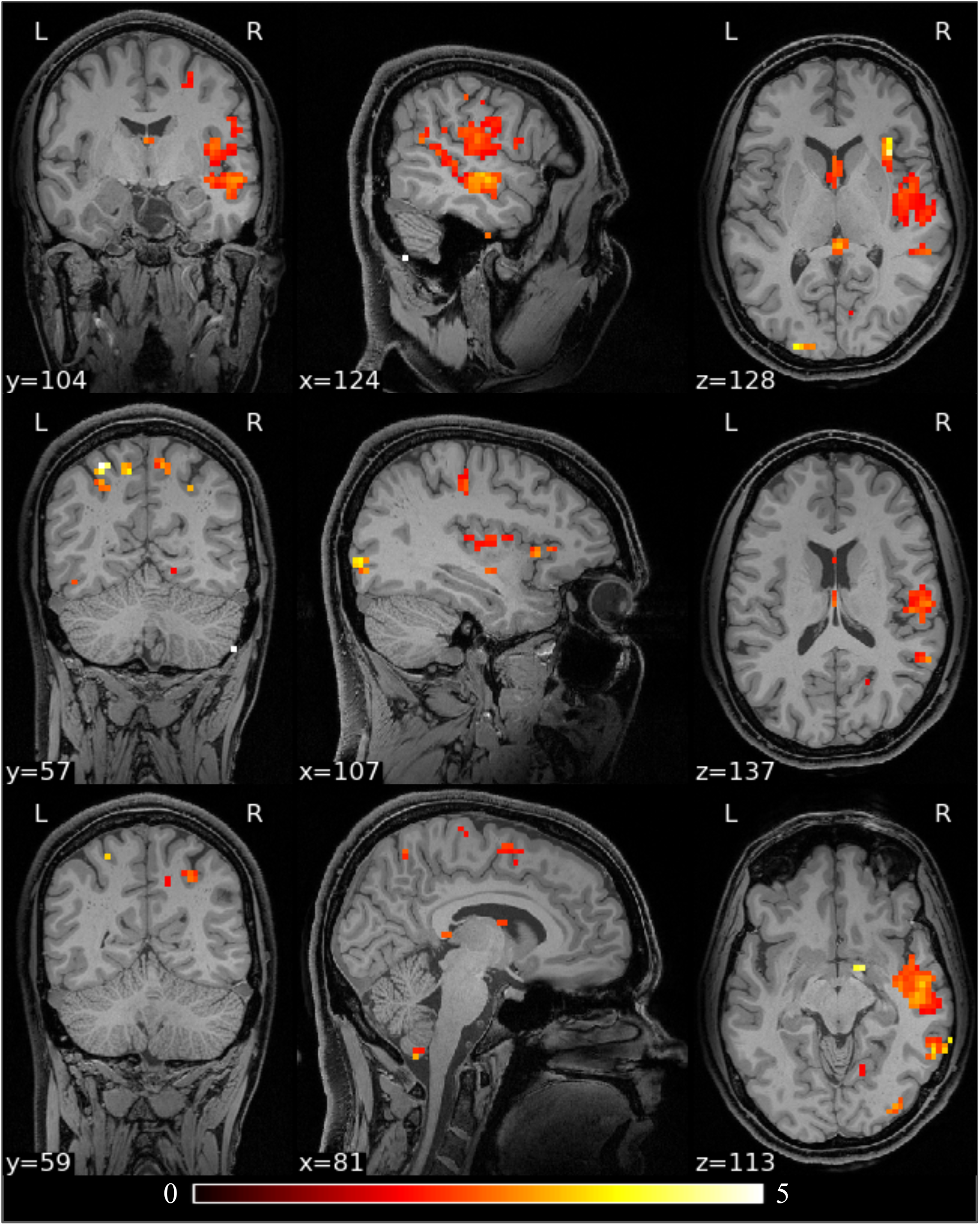
Differences in ALFF between the PTE and non-PTE groups. The results are color-coded f-statistic thresholded by FDR corrected p-values (p < 0.05) derived using a permutation test. Significant clusters are visible in the left temporal lobe, bilateral occipital lobes, cerebellum, and right parietal lobe.

**Figure 3:**
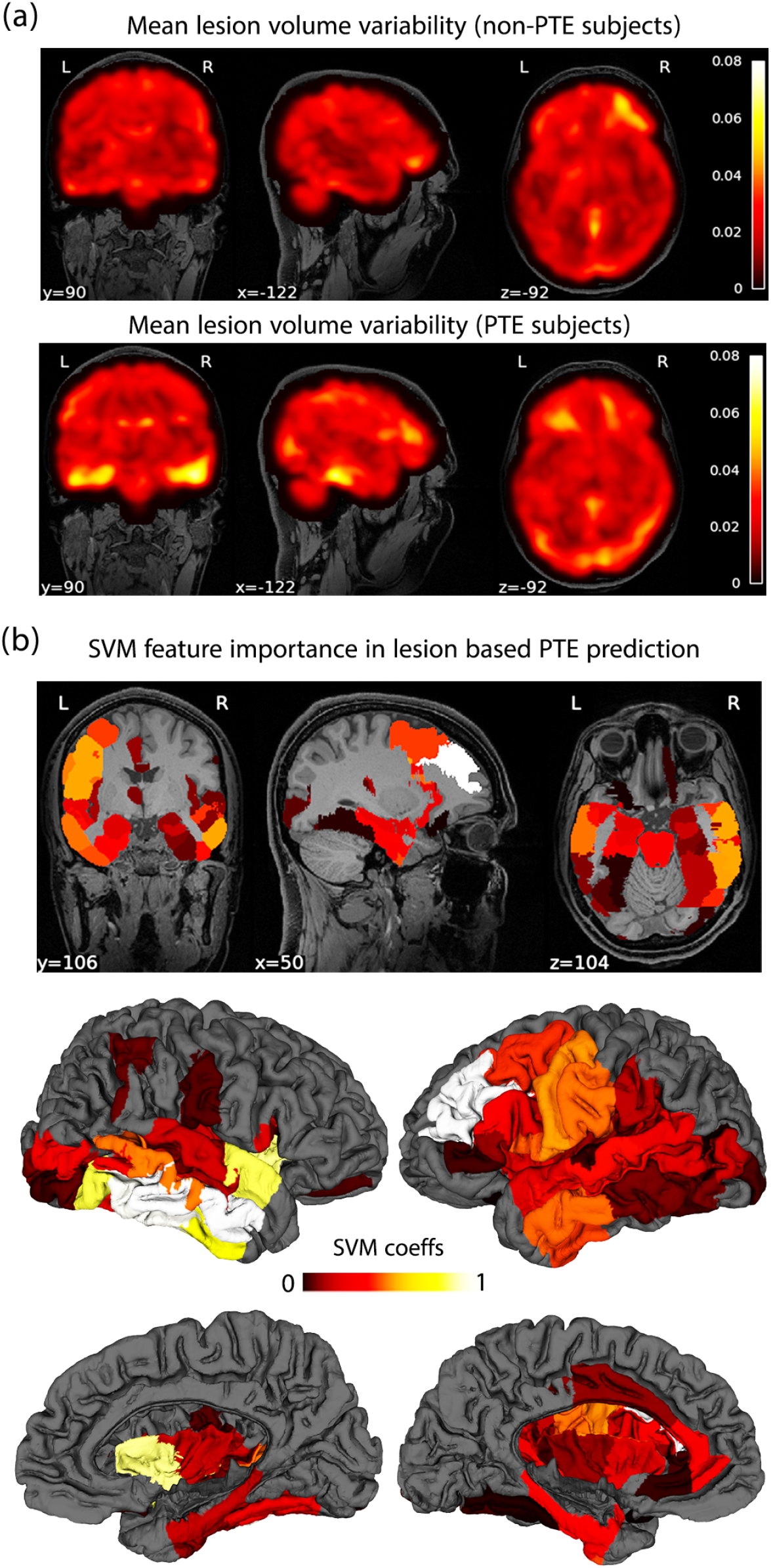
(a) Brain-wide mean lesion volume variability shown for non-PTE (upper row) and PTE subjects (lower row). (b) Feature importance map shown as color-coded ROIs overlaid on the USCBrain atlas. Both corticalsurface and volumetric ROIs are shown.

### 3.2. PTE-related modulations of ALFF

The results of the lobar analysis of ALFF (Table 2) are consistent with the grayordinate-wise analysis (Figure 1), both showing an increased variance in the PTE population relative to non-PTE subjects in the right temporal lobe, both occipital lobes, right parietal lobe. The asymmetry in the lobe-based analysis is possibly due to the limited sample size.

### 3.3. Classification of PTE and non-PTE subjects using Machine Learning

To test the feasibility of training an ML model to distinguish PTE and non-PTE data we performed leave-one-pair-out nested stratified cross-validation over 100 iterations. Nesting was used for parameter tuning (the number of PCA components and model hyperparameter). Single-feature classification using either the lesion, connectivity, or ALFF metrics, was followed by a multi-feature classification approach combining all three features. The input features were computed for distinct ROIs, based on the USCLobes atlas, and were normalized to unit variance, zero mean. From Table 1, we can see that combining all three feature types yields the best model performance in terms of the AUC scores. This is likely a reflection of the complementarity of the information about PTE captured across the lesion, connectivity and ALFF data. Among the four ML methods we used, KSVM achieved the best performance. This is probably due to high variability in the feature space, and improved feature separation through mapping to a higher dimensional space. The neural network performed relatvely poorly on this classification, which given the moderate training sample size, is not surprising.

**Table 1:** PTE vs non-PTE group comparison of lesion and ALFF measures (p-values obtained using an F-test).

| **Lobe** | **P-value (lesion)**<br>**permutation test** | **P-value (ALFF)**<br>**permutation test** |
| --- | --- | --- |
| **Right Temporal** | **0.010** | **0.003** |
| **Left Temporal** | **0.021** | **0.081** |
| **Right Occipital** | **0.031** | **0.035** |
| **Left Occipital** | **0.127** | **0.009** |
| **Right Frontal** | **0.221** | **0.243** |
| **Left Frontal** | **0.326** | **0.177** |
| **Right Parietal** | **0.574** | **0.003** |
| **Left Parietal** | **0.654** | **0.069** |
| **Right Insula** | **0.347** | **0.226** |
| **Left Insula** | **0.546** | **0.724** |
| **Cerebellum** | **0.047** | **0.072** |

**Table 2:** Classification accuracy of PTE vs. non-PTE subjects using different classifiers and features types. Mean and standard deviation of AUC are shown for KSVM, SVM, RF and NN. The last column shows the performance obtained when the models were trained simultaneously on all three feature types.

| Method | Lesion | Connectivity | ALFF | Combined |
| --- | --- | --- | --- | --- |
| KSVM | 0.55(0.04) | 0.61(0.07) | 0.62(0.04) | 0.78(0.04) |
| SVM | 0.63(0.03) | 0.51(0.04) | 0.63(0.02) | 0.64(0.03) |
| RF | 0.58(0.05) | 0.55(0.08) | 0.67(0.03) | 0.64(0.06) |
| NN | 0.50(0.04) | 0.61(0.08) | 0.50(0.05) | 0.56(0.02) |

In addition to determining the feature space based on the USCLobes atlas, we also computed ROI-based features using atlases with a large number of parcels. To this end we used the USCBrain atlas, BCI-DNI atlas [49], and AAL3 atlas [68]. Running the classification pipeline based on these atlases did not lead to any significant improvements in PTE prediction.

In order to gain further insights and improve the interpretability of these ML results, we sought to assess the distinct spatial contributions of the lesion features to the overall prediction score. To this end, we computed feature importance maps derived from the positive SVM model coefficients. Figure 1 shows the lesion variability maps (in non-PTE and PTE subjects), followed by the SVM feature importances across the USCBrain atlas. Comparison between the lesion and feature importance maps points towards a reasonable spatial overlap between the two. But, most importantly, this analysis suggests that the lesion volume data that contribute most to distinguishing PTE and non-PTE subjects are located in right temporal and left prefrontal cortices.

## 4. Discussion

The complex pathophysiology as well as the variability in the degree of severity of TBI poses a significant challenge to any research in this field. TBI with the resultant coup, focal cortical injury, and contrecoup injury, secondary contusion opposite to the coup injury, results in great variety of cortical lesions sites which then gives rise to a diversity of neurocognitive impairments [69]. The cortical injury inflicted by TBI is well known to have some predilection to the polar regions of the brain, particularly the temporal and frontal lobes [70]. This is related to adjacency to bony structures of the skull. However, the presence of cortical lesions in specific regions have not been consistently linked to the development of PTE. In one study a left parietal lobe lesion and presence of hemosiderin staining were linked to the development of PTE [71]. Yet in other studies, rather than a specific cortical region, the degree of leakage by the blood brain barrier around cortical sites after TBI was a prognostic marker for the development of PTE [20]. Due to the extreme heterogeneity of PTE, the need for a reliable biomarker to enhance prediction of PTE is highly desirable. There have been a number of promising and novel therapeutic interventions targeting the complex pathophysiology of TBI shown to have promise in preclinical phase I/II trial, yet have gone on to fail in phase III clinical trials [69]. Some reasons for failure could be the large sample required to determine benefit if the effect size of the intervention is small. Use of biomarkers could enrich the target population so as to maximize the likelihood of discovery of potential therapeutic intervention.

In this study, we explored the feasibility of predicting PTE using functional and structural imaging features which consisted of fMRI connectivity, lesion volumes, and the amplitude of low-frequency fluctuation in FMRI data. In particular, we assessed the performance of widely employed machine learning algorithms, in predicting post-traumatic epilepsy using these features. The aim was two-fold: (1) to assess the feasibility of building predictive ML algorithms for PTE based on functional and structural brain features, (2) leverage this data-driven framework to pinpoint discriminant brain features that may provide useful mechanistic insights into the clinical underpinnings of PTE.

Among the machine learning models examined here, KSVM combined with a standard PCA approach for feature dimensionality reduction led to the highest prediction score. Additionally, our results suggest that combining all three feature types (connectivity, lesion and ALFF data) leads to better prediction than only using one of the three types of features. The feasibility of successfully training a model to discriminate PTE from non-PTE subjects demonstrates that complementary brain features extracted from multi-spectral MRI imaging can collectively capture anatomo-functional alterations that underpin PTE. The ability of the ML approach to generalize to data from individuals that were not used in training the classifier (i.e. using cross-validation) indicates the feasibility of using the identified features to predict with a reasonable degree of accuracy whether a patient who suffers a traumatic brain injury is likely to go on to develop PTE or not. Our results show a maximum AUC of 0.78 using KSVM. Improvements in this value may result from the use of larger training sets as we discuss below.

Although we employed several methods to compare the PTE and non-PTE group, we observed reasonable consistency across the results; The results form the group difference analyses based on F-tests and the ML classifier results, as well as the feature importance maps (based on the SVM coefficients), provide converging evidence for alterations in temporal, and occipital cortices, and to some extent in the cerebellum, in both functional and structural features. We note that the discrepancy between some of the structural and functional patterns observed in the right parietal regions may be due to the functional connectivity between parietal areas and epilepsy-related networks.

Furthermore, it is noteworthy that while the findings from lesion-based analysis were largely left-right symmetric, the ALFF-based analysis results showed a certain degree of asymmetry. To probe this further, we compared lesion volumes in left-hemisphere ROIs and in the corresponding right-hemisphere ROIs. This revealed that the left-right lesion volume differences were not significant for any of the ROIs. In the ALFF-based analysis, the region-wise results were largely symmetric, but in the left temporal, and left parietal lobes, the *p*-values approached significance. The *p*-value in the ALFF analysis for the cerebellum also approached significance.

The AUC metric is a commonly used performance criterion in binary classification. While performance requirements vary across tasks and application domains, an AUC score of 0.7 is frequently used to indicate a minimally acceptable discrimination [72, 73, 74]. However, a higher AUC is often necessary to stratify a significant portion of the population into high-risk or low-risk subjects. Schummers et al. [72] suggested that to classify the majority of the population into a clinically distinct risk group (high or low risk), an AUC of 0.85 was needed. In our analysis, the highest performance (AUC=0.77) was obtained with the KSVM algorithm in a total sample size of 72 individuals (36 PTE and 36 non-PTE subjects). To understand the impact of the number of subjects on the AUC, we repeated the ML analysis for different subsets of subjects (maintaining balanced sample sizes for PTE and non-PTE subjects). Figure 4 depicts the AUC’s mean and variance based on stratified cross-validation as a function of the number of training subjects. Our results suggest that the AUC starts to increase monotonically after 26 subjects. A quadratically fit curve that extrapolates this trend is shown as a dotted red line. We fit a curve to the last 6 data points to obtain an over-optimistic estimate of 0.98 AUC for 50 PTE subjects. By fitting the last 10 points we obtain a more conservative AUC value of 0.85 for 50 PTE subjects. Assuming a 15% prevalence of PTE in TBI subjects, this analysis shows that N=300 would yield the desired AUC with the existing algorithm. This is, of course, an attempt to extrapolate our results to outline a pragmatic and clinically relevant approach to using our method for the prediction of PTE. Of course, there are a number of unknown variables and factors that can influence this analysis. These include the heterogeneity of TBI and a wide range of properties of TBI patients (incl. demographics, types, the severity of injuries, treatment at the acute stage, etc.). In particular, the properties of the Maryland data used for this analysis can be quite different from those of other data sets.

**Figure 4:**
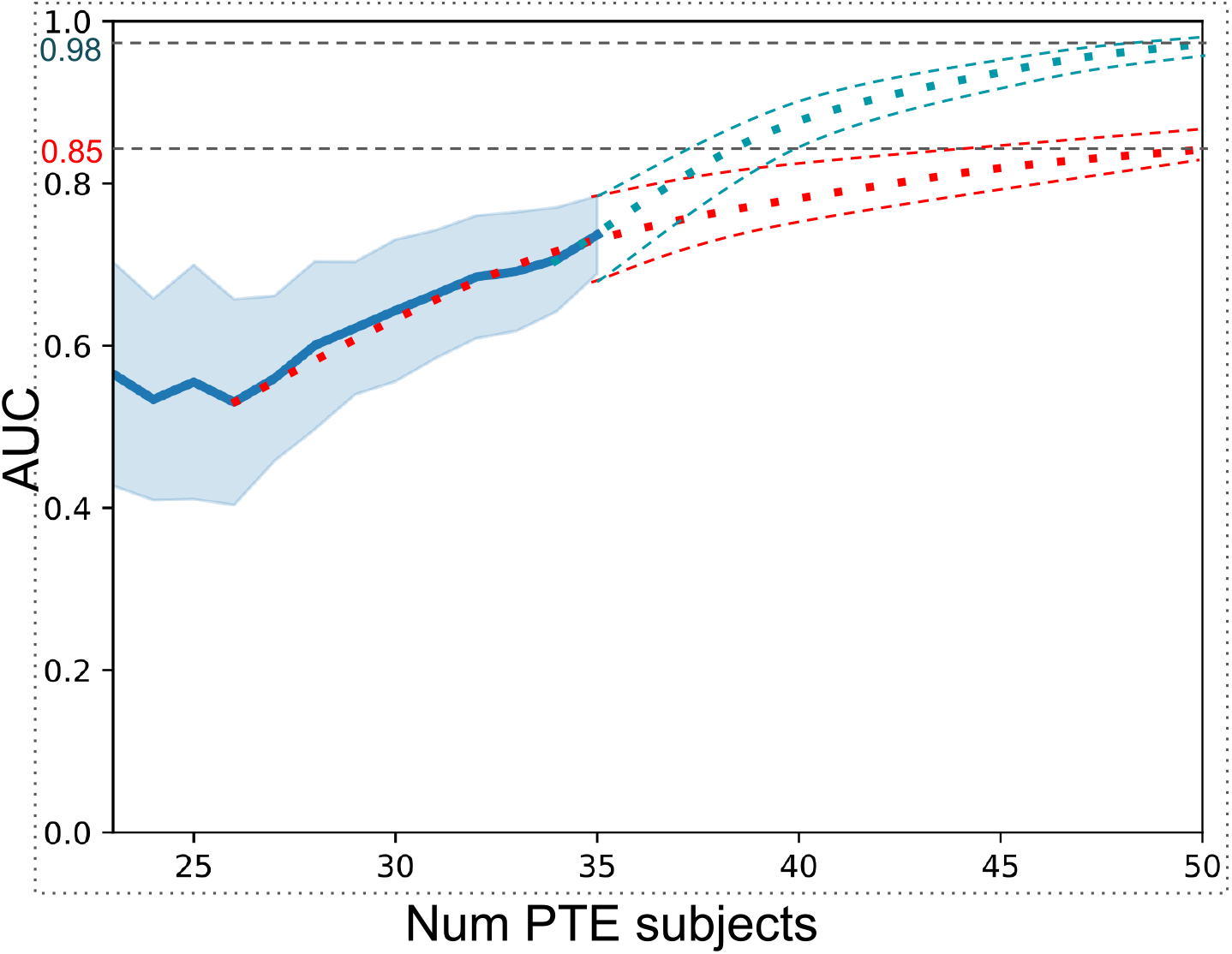
Number of samples vs AUC resulting from KSVM (PCA) method. The blue curve shows mean AUC and shaded areas indicate std dev. in the leave one out stratified cross-validation. Conservative (red) and optimistic (blue) extrapolations are shown as dotted curves.

### 4.1. Limitations

Despite the encouraging and clinically relevant observations reported here, the PTE prediction method proposed is a first step that paves the way for more efficient and elaborate prediction approaches. Among relevant future steps, we believe that increasing the size of the training data and the addition of more features such as diffusion imaging data may lead to an improved AUC. Similarly, prediction is likely to benefit from incorporating non-imaging clinical data such as scores on the Glasgow Coma Scale (GCS), which classifies Traumatic Brain Injuries, alongside demographics, and injury mechanisms. As noted earlier, this information was unfortunately not available in the public domain for the cohort we used in this study. Furthermore, including information on the type of epilepsy, may turn out to be useful for training the ML models. This and other clinical information can also be used for further sub-grouping of the clinical population.

## 5. Conclusion

In this paper, we investigated the efficiency of functional and structural brain features as PTE biomarkers. Leveraging a machine-learning framework, we compared PTE prediction performance across an array of standard classifiers and a variety of brain features. The best results were obtained with KSVM, which is possibly partly due to the heterogeneity of the alterations in the PTE group around the mean feature. Our results using kernel-based methods show promising results. In both lesion and ALFF comparison studies, bilateral temporal lobes and cerebellum show significance. Moreover, there is potential involvement of the parietal and occipital lobes. Cross-sectional studies of children with chronic localization-related epilepsy (LRE) using traditional volumetric and voxel-based morphometry have revealed abnormalities in the cerebellum, frontal and temporal lobes, hippocampus, amygdala, and thalamus [75, 76, 77, 78, 79, 80, 81]. The temporal lobe findings from our study further support this evidence.

One of the limitations of our study is its relatively small population size (N=74). A larger study using TrackTBI dataset might be possible in the future. The leave-one-out cross-validation shows the relatively stable performance of the prediction. However, the AUC performance can be improved further with a larger dataset.

